# Lateralised Transmission of Beta Oscillations From Cortex to Muscle During Movement Cancellation Reflects Selective Motor Inhibition

**DOI:** 10.64898/2026.09.11.750861

**Authors:** Ciarán McGeady, Jaime Ibáñez Pereda, María Sarasquete, Tianyu Jia, Dario Farina

**Affiliations:** Department of Bioengineering, Imperial College London, London, United Kingdom; BSICoS, I3A and IIS Aragón, University of Zaragoza, Zaragoza, Spain; Centro de Investigación Biomédica en Red en Bioingeniería, Biomateriales y, Nanomedicina (CIBER-BBN), Zaragoza, Spain

**Keywords:** electrophysiology, motor control, motor inhibition, movement cancellation

## Abstract

Beta oscillations in the sensorimotor cortex are modulated across various stages of motor control, including movement preparation and cancellation, and are known to project to the motor neuron pool and muscles during Go/No-Go tasks. While these oscillations have been linked to motor inhibition, the specificity of their transmission to end-effector muscles during movement cancellation remains unclear. We investigated the selectivity of beta-band activity using a lateralised Go/No-Go paradigm. Participants maintained steady bimanual wrist extensor contractions and either executed a unilateral ballistic wrist extension (Go) or cancelled the planned movement (No-Go). Concurrent electroencephalographic and electromyographic recordings were measured throughout. Successful movement cancellation was characterised by contralateral cortical beta modulation accompanied by stronger and more clearly lateralised peripheral beta synchronisation in the muscle associated with the suppressed movement. Peripheral beta activity exhibited greater magnitude and clearer separability between left- and right-sided conditions than cortical signals. These findings demonstrate effector-specific brain-to-muscle beta transmission during movement cancellation and highlight the potential of peripheral measures to provide enhanced resolution of motor cortical dynamics.

## Introduction

Beta-band oscillations (13–30 Hz) are a prominent characteristic of sensorimotor cortical activity and are closely associated with the preparation, execution, and suppression of voluntary movement [1]. In Go/No-Go paradigms, movement cancellation is consistently accompanied by modulation of cortical beta activity, implicating beta-band dynamics in the neural control of motor inhibition [2], [3], [4]. Interestingly, these oscillations are not confined to the brain: Cortical beta signals propagate along the corticospinal tract and are coupled to the activity of spinal motor pools and peripheral musculature [5], [6], [7]. Beta oscillations therefore may provide a link between cortical processes underlying motor inhibition and their expression throughout the descending motor system. However, whether inhibitory beta dynamics associated with movement cancellation are transmitted to the periphery, and how this transmission is organised across muscles, remains poorly understood.

Zicher et al. [8] provided initial evidence that No-Go beta modulation extends beyond the sensorimotor cortex to a peripheral muscle (tibialis anterior). However, examination of a single muscle leaves unresolved whether inhibitory beta activity is broadcast broadly across the active motor pool or is selectively transmitted to the representation of the effector being suppressed. Distinguishing between these possibilities is critical for understanding how cortical inhibitory commands are distributed through the descending motor system.

Two competing models make distinct predictions. A global inhibition framework, supported by behavioural and physiological evidence, predicts that cancellation engages a non-selective inhibitory signal across the motor system. For example, stopping one hand during a bimanual task delays execution of the other hand by approximately 100 ms [9], while motor-evoked potentials are suppressed in task-irrelevant muscles during unilateral movement cancellation [10]. If beta oscillations constitute a component of this global inhibitory signal, their propagation should similarly extend across active motor representations. Alternatively, inhibitory control may be implemented through effector-specific channels, in which case downstream beta transmission should be expressed within the cortical and peripheral representations of the cancelled effector. These contrasting predictions provide a means of testing whether beta-band communication during movement cancellation reflects a global inhibitory broadcast or spatially targeted control.

Here, we address this gap using a bimanual Go/No-Go paradigm with simultaneous electroencephalography (EEG) and bilateral electromyography (EMG). Participants maintained steady bilateral wrist extensor contractions while responding to lateralised visual cues, withholding a planned unimanual extension on No-Go trials while keeping baseline bilateral force constant. By applying linear mixed-effects models to trial-level time–frequency data, we show that post-No-Go related beta modulation is lateralised to the contralateral cortex and that peripheral beta modulation displays even greater effector specificity and magnitude than its cortical counterpart. These findings demonstrate spatially targeted corticospinal beta transmission during movement cancellation, uncovering effector-specific beta modulation at the periphery. A preliminary version of this work was previously presented in abstract form at [11].

## Results

### Behavioural data

Force recordings from a representative participant show the contrast between the Go and No-Go conditions (Fig. 1a). Following an initial force ramp to a 10% MVC baseline plateau, Go trials produced a rapid force exertion restricted to the cued wrist (black traces). In contrast, during No-Go trials, force output in the targeted wrist was maintained at the 10% MVC plateau. All trials concluded with a force decline back to 0% MVC. During Go trials, reaction times for cued unilateral wrist extensions did not significantly differ between left (410.5 *±* SD ms) and right (418.8 *±* SD ms) sides (*p* = 0.51; Fig. 1b).

**Figure 1.**
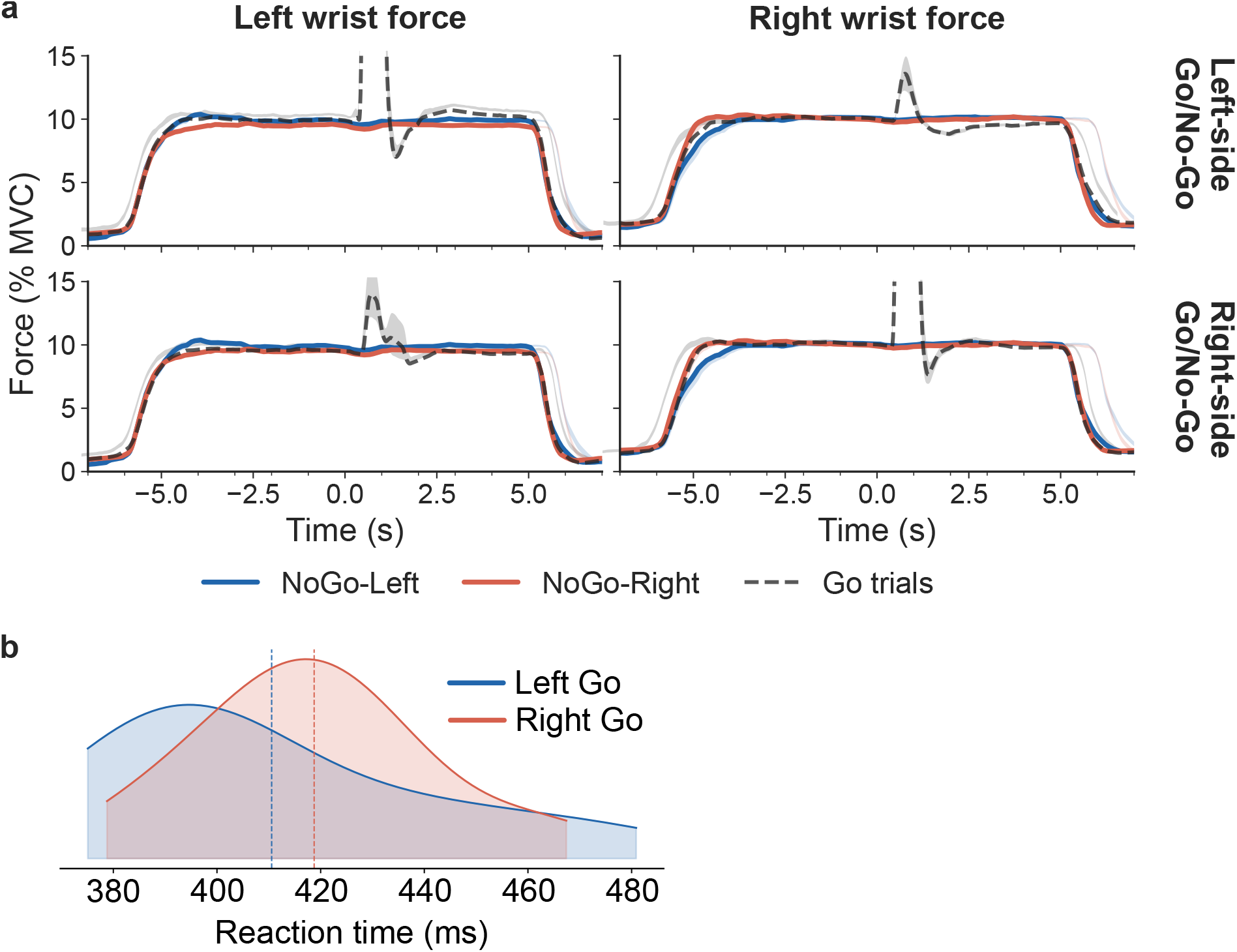
Behavioural data. **a**, Wrist force traces for left- and right-sided movement cancellation conditions in a representative participant. Each 14 s trial began at 0% MVC, followed a ramp to a bilateral 10% MVC plateau, with the Go/No-Go cue at 0 s. Top: left-wrist cancellation; bottom: right-wrist cancellation. Go trials are shown in black. **b**, Reaction times in response to the Go cue for left- and right-sided Go trials.

### Movement cancellation elicits contralateral cortical beta modulation

The temporal profile of cortical beta power across conditions is shown in Fig. 2 and Fig. 3. The linear mixed model revealed significant main effects of Hemisphere (*χ*^2^(6) = 54.87, *p* = 4.92 *×* 10^−10^), Condition (*χ*^2^(3) = 19.67, *p* = 0.0002) and Time Window (*χ*^2^(2) = 95.68, *p* = 1.67 *×* 10^−21^, 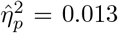), as well as significant two-way interactions of Hemisphere *×* Condition (*χ*^2^(3) = 35.41, *p <* .001), Hemisphere *×* Window (*χ*^2^(4) = 46.93, *p <* .001) and Condition *×* Window (*χ*^2^(4) = 33.73, *p <* .001). All effects were guided by a significant three-way interaction of Hemisphere, Condition and Time Window (*χ*^2^(2) = 32.85, *p* = 7.37 *×* 10^−8^), indicating that the temporal profile of beta modulation differed between hemispheres as a function of which wrist movement was being inhibited.

**Figure 2.**
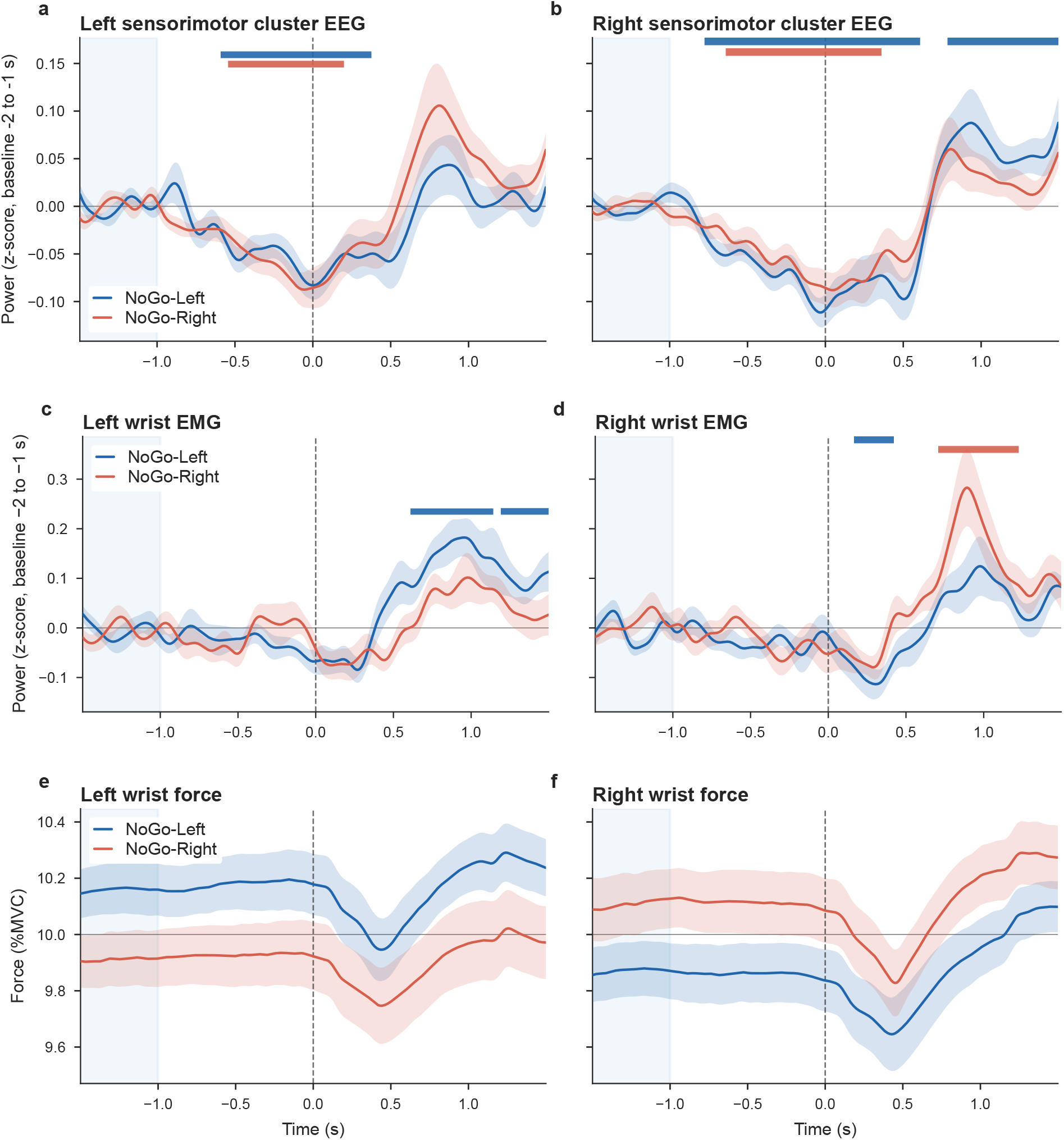
Time courses of cortical and peripheral beta-band power during movement cancellation. Beta-band power (*z*-scored relative to a −2 to −1 s baseline) for NoGo-Left (blue) and NoGo-Right (red) conditions, over time relative to the No-Go cue (dashed vertical line at 0 s). Shaded regions denote *±* 1 s.e.m. across subjects. The shaded blue rectangle indicates the baseline period. **a**, Left sensorimotor cluster. **b**, Right sensorimotor cluster. **c**, Left wrist EMG. **d**, Right wrist EMG. Horizontal bars indicate periods of statistically significant beta modulation relative to baseline, as determined by cluster-based permutation testing (*p <* 0.05; blue: NoGo-Left; red: NoGo-Right). **e, f** show wrist extensor force.

**Figure 3.**
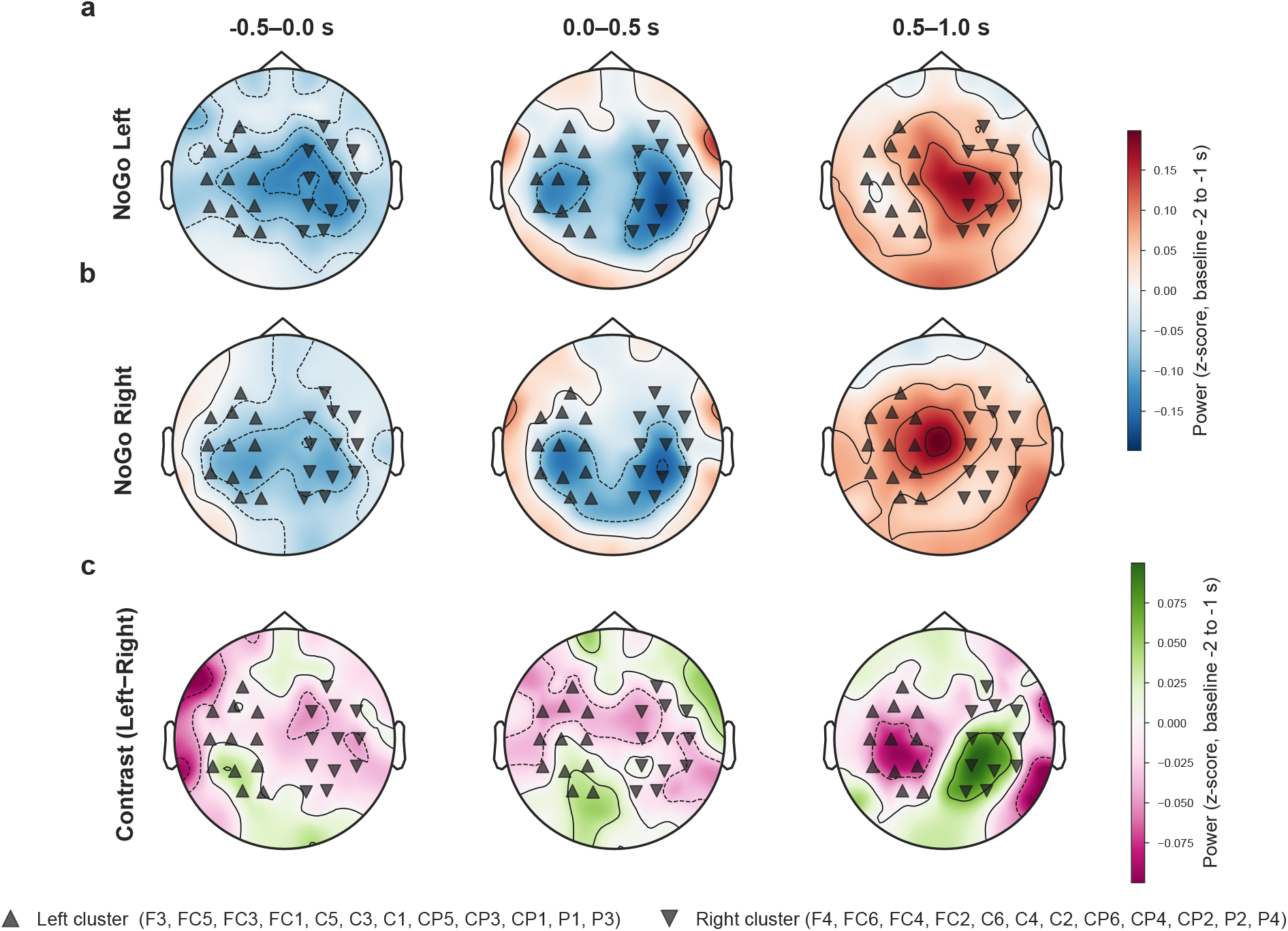
Topographic distribution of cortical beta-band power during movement cancellation. Average scalp topographies beta-band power during movement cancellation (*z*-score normalised relative to a −2 to −1 s baseline) across participants across three time windows: pre-No-Go (−0.5–0.0 s), early No-Go (0.0–0.5 s), and late No-Go (0.5–1.0 s). **a**, NoGo-Left trials. **b**, NoGo-Right trials. **c**, Condition contrast (NoGo-Left minus NoGo-Right). Black triangular markers denote the left and right sensorimotor electrode clusters used for analysis.

### Pre-cancellation window (−0.5–0 s)

Beta power was significantly suppressed below baseline across both hemispheres and both conditions in the pre-stimulus window (means: −0.055 to −0.079; all FDR-corrected *p* ≤ 0.003, |*d*| = 1.05–1.62), Fig. 3a, Fig. 4a, Fig. **??**a. The largest suppression was observed in the right sensorimotor cluster on NoGo-Left trials (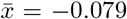 ; *t*(14) = −6.26, *p*_FDR_ = 0.0003, *d* = −1.62), followed by the right sensorimotor cluster on NoGo-Right trials (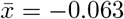 ; *t*(14) = −4.47, *p*_FDR_ = 0.003, *d* = −1.15), the left sensorimotor cluster on NoGo-Right trials (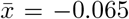 ; *t*(14) = −4.35, *p*_FDR_ = 0.003, *d* = −1.12), and the left sensorimotor cluster on NoGo-Left trials (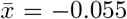 ; *t*(14) = −4.08, *p*_FDR_ = 0.003, *d* = −1.05). This bilateral, condition-independent suppression is consistent with anticipatory motor cortical desynchronisation reflecting a non-specific state of heightened motor readiness prior to the No-Go stimulus, and also resembles the contingent negative variation signal.

**Figure 4.**
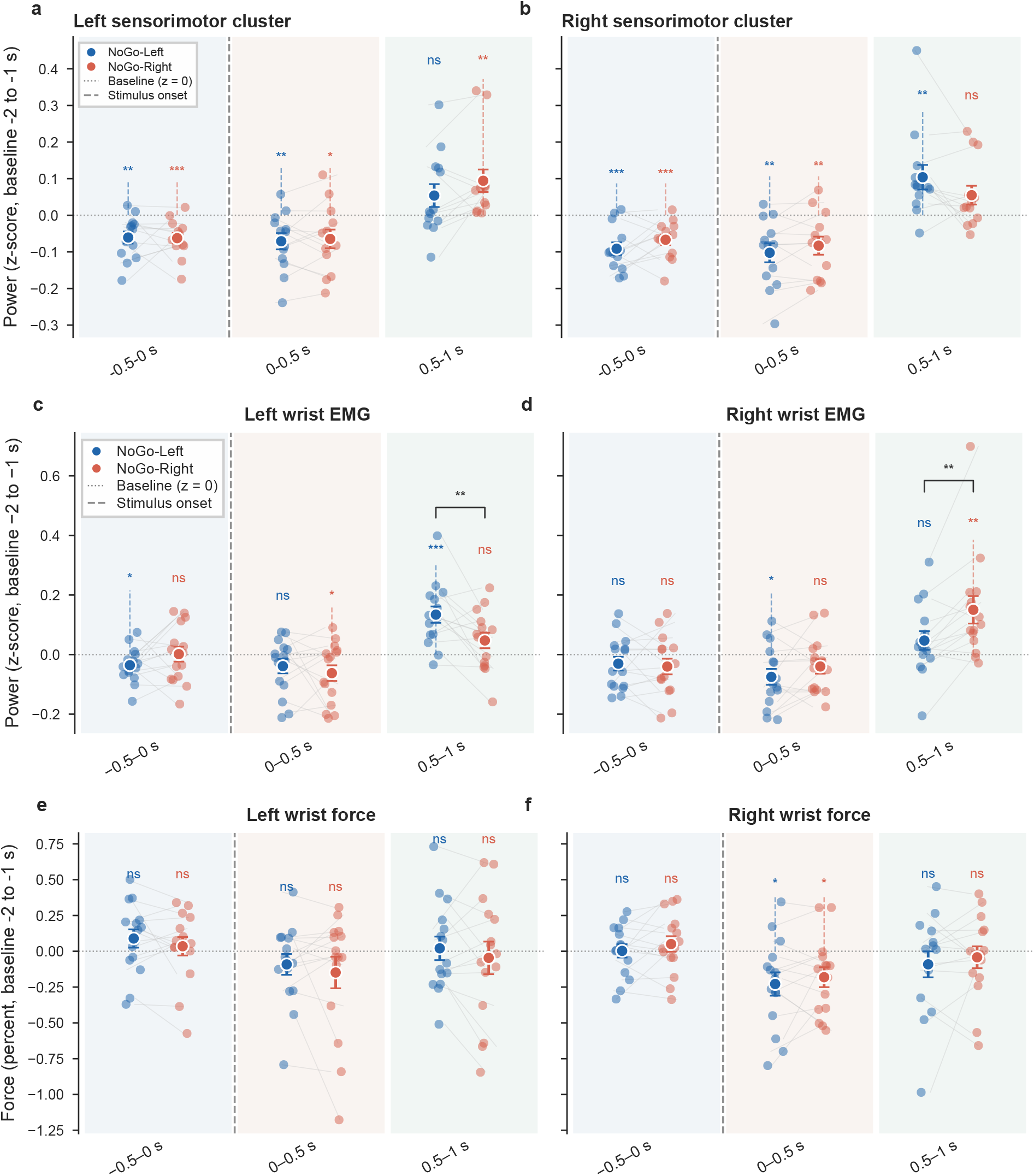
Window-averaged beta power and force during movement cancellation. Subject-level data for NoGo-Left (blue) and NoGo-Right (red) conditions across three analysis windows: pre-cancellation (−0.5–0.0 s), early cancellation (0.0–0.5 s), and late cancellation (0.5–1.0 s). Large filled circles denote group means *±* s.e.m.; small circles represent individual subjects, connected by grey lines. The horizontal dotted line indicates baseline (*z* = 0). **a**, Left sensorimotor cluster. **b**, Right sensorimotor cluster. **c**, Left wrist EMG. **d**, Right wrist EMG. **e**, Left wrist force. **f**, Right wrist force. Beta power (*z*-scored relative to a −2 to −1 s baseline) is shown in panels **a**–**d**; force (percentage change from baseline) is shown in panels **e**–**f.** Asterisks above individual condition data denote significant modulation from baseline (one-sample *t*-tests, FDR-corrected; ^∗^*p <* 0.05, ^∗∗^*p <* 0.01, ^∗∗∗^*p <* 0.001). Bracketed asterisks between conditions denote significant pairwise differences (paired *t*-tests, Holm-corrected). ns, not significant.

**Figure 5.**
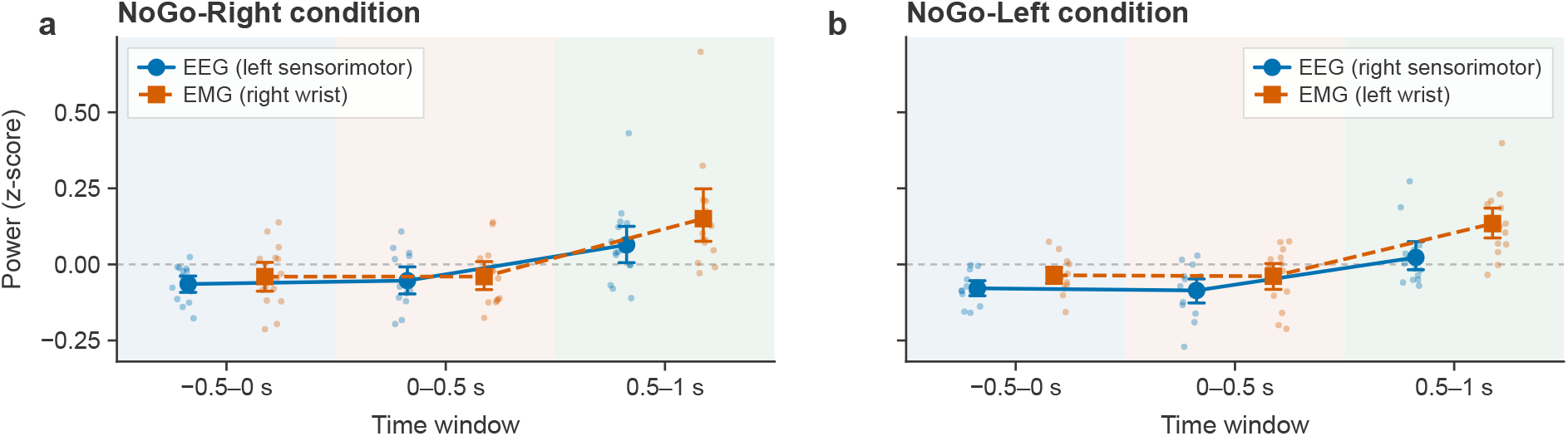
Contralateral comparison. Window-averaged beta power (*z*-scored relative to a −2 to −1 s baseline) for the contralateral EEG sensorimotor cluster (blue) and ipsilateral wrist EMG (red) across three time windows: pre-No-Go (−0.5–0.0 s), early No-Go (0.0–0.5 s), and late No-Go (0.5– 1.0 s). Large dots denote group means *±* s.e.m.; small dots represent individual subjects. **a**, NoGo-Right condition: left sensorimotor EEG cluster versus right wrist EMG. **b**, NoGo-Left condition: right sensorimotor EEG cluster versus left wrist EMG. In both conditions, peripheral beta modulation in the late cancellation window exceeds the magnitude of the cortical response, suggesting that effector-specific beta synchronisation following movement cancellation is more pronounced at the peripheral than at the cortical level.

### Early cancellation window (0–0.5 s)

Beta suppression persisted into the early post-stimulus window across all four cells (means: −0.053 to −0.086), with all cells surviving FDR correction. The deepest suppression was observed in the right sensorimotor cluster on NoGo-Left trials (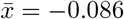 ; *t*(14) = −4.10, *p*_FDR_ = .003, *d* = −1.06), followed by the right sensorimotor cluster on NoGo-Right trials (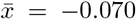 ; *t*(14) = −3.81, *p*_FDR_ = .004, *d* = −0.98), the left sensorimotor cluster on NoGo-Left trials (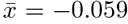 ; *t*(14) = −3.14, *p*_FDR_ = .012, *d* = −0.81), and the left sensorimotor cluster on NoGo-Right trials (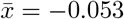 ; *t*(14) = −2.37, *p*_FDR_ = .039, *d* = −0.61). The bilateral, condition-independent nature of this suppression is consistent with a non-lateralised inhibitory control mechanism engaging both sensorimotor hemispheres symmetrically during the active stopping period.

### Late cancellation window (0.5–1 s)

In the late post-stimulus window, beta power rebounded above baseline, and the pattern of FDR-corrected significance dissociated cleanly along contralateral versus ipsilateral lines (Fig. 4. Indeed this can also be seen in the topographical contrast in Fig. 3 c. Both contralateral cells survived FDR correction: the right sensorimotor cluster on NoGo-Left trials (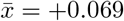 ; *t*(14) = 2.52, *p*_FDR_ = 0.037, *d* = +0.65) and the left sensorimotor cluster on NoGo-Right trials (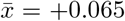 ; *t*(14) = 2.40, *p*_FDR_ = .039, *d* = +0.62). By contrast, neither ipsilateral cell reached significance after correction: the left sensorimotor cluster on NoGo-Left trials showed a negligible rebound (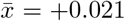 ; *t*(14) = 0.80, *p*_FDR_ = .439, *d* = +0.21), and the right sensorimotor cluster on NoGo-Right trials showed a small, non-significant rebound 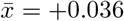 *t*(14) = 1.78, *p*_FDR_ = .107, *d* = +0.46). This contralateral selectivity was confirmed by the three-way interaction term for the right hemisphere NoGo-Right late-window contrast (*β* = −0.095, SE = 0.018, *z* = −5.23, *p <* .001).

These results indicate a two-phase pattern of cortical beta modulation during No-Go inhibition: an early, bilateral desynchronisation spanning the pre-No-Go and post-No-Go periods, followed by a spatially selective synchronisation contralateral to the suppressed response hand. This dissociation between the non-specific ERD and the lateralised ERS suggests that while motor preparation and active inhibition engage both hemispheres symmetrically, the post-inhibitory beta rebound reflects a unilateral motor reset signal that is tuned to the effector being suppressed (important). We consider whether this pattern extends beyond the cortical region to the peripheral level in the next section.

### Movement cancellation elicits ipsilateral peripheral beta modulation

The temporal profile of peripheral beta power across conditions is shown in Figure 2c and Figure 2d. The LMM revealed significant main effects of Channel (*χ*^2^(6) = 53.09, *p <* .001), Condition (*χ*^2^(3) = 23.16, *p <* .001) and Time Window (*χ*^2^(2) = 39.20, *p <* .001), as well as significant two-way interactions of Channel *×* Condition (*χ*^2^(3) = 51.59, *p <* .001), Channel *×* Window (*χ*^2^(4) = 34.90, *p <* .001) and Condition *×* Window (*χ*^2^(4) = 34.59, *p <* .001). All effects were subsumed by a significant three-way interaction of Channel, Condition and Time Window (*χ*^2^(2) = 34.25, *p <* .001), indicating that the temporal profile of beta modulation in wrist EMG differed between channels as a function of which movement was being cancelled. The random slope structure was justified by a likelihood ratio test against a random-intercept-only model (*χ*^2^(2) = 64.55, *p <* .001). One subject (Subject 4) was identified as a univariate outlier in the right wrist EMG NoGo-Right late-window cell (*z* = 3.07); all reported results are based on the full sample (*N* = 15), and conclusions were unchanged when this subject was excluded.

### Pre-cancellation window (−0.5–0 s)

Beta power was not significantly different from baseline in any cell in the pre-stimulus window (all FDR-corrected *p* ≥ 0.080, |*d*| ≤ 0.58), and no significant condition differences were observed in either channel (both *p*_Holm_ ≥ .727). Although the left wrist EMG showed a numerical trend toward suppression on NoGo-Left trials (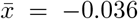 ; *t*(14) = −2.40, *p*_FDR_ = .080, *d* = −0.62), this did not survive correction. This bilateral, condition-independent pattern indicates that prior to stimulus onset, beta power was symmetrically distributed across both wrist EMG channels regardless of which movement would subsequently be cancelled, consistent with a non-specific state of bilateral isometric contraction.

### Early cancellation window (0–0.5 s)

In the early post-stimulus window, beta power remained numerically suppressed below baseline across all four cells, though no cell survived FDR correction (all *p*_FDR_ ≥ .058). The deepest suppression was observed in the right wrist EMG on NoGo-Left trials (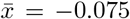 ; *t*(14) = −2.79, *p*_FDR_ = .058, *d* = −0.72), followed by the left wrist EMG on NoGo-Right trials (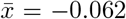 ; *t*(14) = −2.36, *p*_FDR_ = .080, *d* = −0.61). Condition contrasts were non-significant in both channels (both *p*_Holm_ ≥ .727). The broadly symmetric, condition-independent suppression at this window is consistent with a non-lateralised inhibitory mechanism engaging both wrist representations during the active stopping period.

### Late cancellation window (0.5–1 s)

In the late post-No-Go window, beta power rebounded above baseline, and the pattern dissociated sharply along ipsilateral versus contralateral lines. Both ipsilateral cells survived correction: the left wrist EMG on NoGo-Left trials (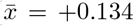 ; *t*(14) = 4.89, *p*_FDR_ = 0.003, *d* = +1.26) and the right wrist EMG on NoGo-Right trials (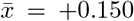 ; *t*(14) = 3.26, *p*_FDR_ = 0.034, *d* = +0.84). By contrast, neither contralateral cell reached significance: the right wrist EMG on NoGo-Left trials showed only a small, non-significant rebound (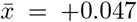 ; *t*(14) = 1.50, *p*_FDR_ = 0.186, *d* = +0.39), and the left wrist EMG on NoGo-Right trials showed a comparable non-significant rebound (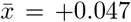 ; *t*(14) = 1.83, *p*_FDR_ = .179, *d* = +0.47). This ipsilateral selectivity was confirmed by direct condition contrasts: the left wrist EMG showed significantly greater beta power during NoGo-Left than NoGo-Right trials (mean difference = +0.087; *t*(14) = 3.02, *p*_Holm_ = .046, *d* = +0.78), while the right wrist EMG showed the opposite pattern, with significantly greater beta power during NoGo-Right than NoGo-Left trials (mean difference = −0.103; *t*(14) = −3.65, *p*_Holm_ = .016, *d* = −0.94). These opposing condition contrasts across channels indicate a lateralised effect and are reflected in the three-way interaction coefficient for the right wrist EMG NoGo-Right late-window contrast (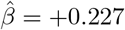, SE = 0.039, *z* = 5.83, *p <* .001).

These results demonstrate that beta power modulation in wrist EMG is lateralised as a consequence of the effector being cancelled. An early, broadly bilateral suppression spanning the pre-stimulus and inhibition-period windows (all *p*_FDR_ ≥ .058, |*d*| ≤ 0.72) is consistent with non-lateralised motor preparation and active stopping engaging both wrist representations symmetrically. This is followed by a spatially selective post-inhibitory beta rebound, significant only in the EMG channel ipsilateral to the cancelled movement (*d* = +0.84–+1.26, both *p*_FDR_ ≤ .034), while the contralateral channel fails to rebound significantly (*d* = +0.39–+0.47, both *p*_FDR_ ≥ .179). The large-to-medium effect sizes of the ipsilateral rebound, combined with the significant opposing condition contrasts in each channel (*p*_Holm_ ≤ .046), indicate that the peripheral muscular signature of movement cancellation includes a lateralised beta band synchronisation that is tuned to the effector representation of the suppressed response limb.

### Force does not account for beta lateralisation

Force was measured in parallel with EEG and EMG (Fig.2 e,f). To assess whether the lateralised peripheral beta rebound could be attributed to differences in wrist force output between conditions, we conducted three complementary analyses using wrist force recorded in parallel with EMG.

First, an LMM of identical structure applied to force revealed a significant Channel *×* Condition interaction (*χ*^2^(3) = 119.55, *p <* .001), confirming that force was itself lateralised: the left wrist showed greater force on NoGo-Left than NoGo-Right trials, and the right wrist showed the opposite pattern (all *p*_Holm_ ≤ 0.048, all |*d*| ≤ 0.32), suggesting participants were pressing slightly harder with the ipsilateral wrist to the cancelled movement, as illustrated in Fig. 2 e, f. However, critically, the Channel *×* Condition *×* Window interaction was entirely non-significant (*χ*^2^(2) = 0.36, *p* = .835), indicating that force lateralisation was temporally invariant, present equally across the pre-No-Go, and post-No-Go windows. By contrast, beta lateralisation emerged *specifically* in the late cancellation window (0.5–1 s). This temporal dissociation between a time-invariant force lateralisation and a time-specific beta lateralisation constitutes strong evidence against force as a mechanistic explanation for the beta effect.

Second, including centred force as a covariate in the beta LMM significantly improved model fit (*χ*^2^(1) = 18.70, *p <* .001), with a positive force–beta coefficient (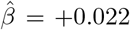, SE = 0.005, *z* = 4.32, *p <* .001), indicating that higher force was associated with higher beta power. However, the critical Channel *×* Condition *×* Window interaction remained significant and unchanged after force adjustment (*χ*^2^(2) = 40.20, *p <* .001; cf. uncontrolled: *χ*^2^(2) = 34.25, *p <* .001), confirming that the lateralised beta rebound was not accounted for by force.

Third, within-subject trial-level Pearson correlations between force and beta power were non-significant across all Channel *×* Condition *×* Window cells after Benjamini–Hochberg correction (all *p*_FDR_ ≥ .387, all |*r*| ≤ .13, all 95 % CIs spanning zero), indicating that the between-subject force–beta association did not reflect a within-subject trial-level coupling.

These analyses reveal a double dissociation: force lateralisation was present but temporally invariant across all windows, whereas beta lateralisation was temporally specific to the post-NoGo period. The positive force–beta association reflected individual differences in overall motor output level rather than a trial-level mechanistic link. The lateralised beta rebound therefore represents a neural signature of effector-specific movement cancellation that is dissociable from the peripheral force profile of the suppressed response, both in its temporal dynamics and at the level of individual trials.

## Discussion

This study investigated the spatiotemporal profile of beta oscillatory power (15–35 Hz) during movement cancellation in a bimanual Go/No-Go paradigm, examining both cortical and peripheral muscular activity simultaneously to determine whether beta modulation is transmitted selectively to the effector representation of the suppressed limb or broadcast non-specifically across all active motor representations. Participants maintained steady bimanual wrist extensor force while preparing a unimanual ballistic contraction in response to an unpredictable Go cue. On No-Go trials, the planned movement was cancelled and bilateral force was sustained. Linear mixed models applied to trial-level beta power revealed contralateral increases in cortical power alongside stronger, lateralised peripheral beta modulation in the muscle corresponding to the cancelled movement. These findings indicate that movement cancellation is followed by effector-specific cortical–peripheral transmission, supporting a targeted rather than globally distributed inhibitory mechanism.

Our present findings extend prior work by Zicher et al. [8], who demonstrated cortical-to-muscle beta transmission to the tibialis anterior in Go/No-Go, and are consistent with a growing body of studies on beta bursting and corticomuscular coherence showing that beta oscillations propagate beyond cortical structures to spinal motor neurons and peripheral muscles [12], [13], [14], [15]. While many studies report findings on the nature of beta oscillatory activity, beta is likely not a single centrally generated signal and must be taken in context. Before contextualising these findings within the broader inhibitory control literature, it is important to consider how the specific features of the present experimental protocol may have shaped the observed beta dynamics [16], [17]. The paradigm employed a bimanual design in which participants were required to cancel one hand’s movement while executing the other’s, with the No-Go cue appearing at a fixed, predictable interval following the Go signal. This design departs from the canonical Go/No-Go paradigm [18] in several important respects. First, the requirement to simultaneously execute and cancel movements in opposite hands introduces a degree of inter-limb coordination that is absent in unimanual stopping tasks, and may engage additional cortical and subcortical circuits involved in bimanual coordination [19]. Second, and perhaps more importantly, the predictability of the No-Go cue means that participants could anticipate its arrival and prepare their potential inhibitory response in advance, rather than reacting to an unexpected No-Go stimulus. This distinction between *proactive* and *reactive* inhibitory control is important for interpreting the present results and is discussed in detail below.

The modulation of sensorimotor beta power during response inhibition has been strongly linked to the hyperdirect pathway connecting prefrontal cortex directly to the subthalamic nucleus (STN), bypassing the striatum [4], [20]. Under this framework, a broad inhibitory signal is broadcast from the right inferior frontal cortex (rIFC) via the hyperdirect pathway to the STN, which in turn suppresses thalamo-cortical drive and reinstates beta synchrony across sensorimotor networks [21]. Consistent with this model, beta power increases following successful stopping have been reported in the STN and sensorimotor cortex [22]. This basal ganglia–thalamocortical circuit can, in principle, produce two different patterns of beta modulation depending on the specificity of the inhibitory signal: a global pattern, in which beta synchronisation is broadcast non-specifically to all active motor representations regardless of which effector is being cancelled, or a *specific* pattern, in which the inhibitory signal is targeted selectively to the motor representation of the to-be-cancelled effector [9], [23]. The present results support the specific modulation hypothesis since the post-No-Go beta rebound was lateralised at both the cortical and peripheral levels, with the largest rebounds observed in the hemisphere and muscle contralateral and ipsilateral to the cancelled movement, respectively. This is consistent with reports of effector-specific beta modulation during selective stopping in tasks requiring partial response inhibition [9], [23], and argues against a purely global, non-specific inhibitory broadcast as the mechanism underlying movement cancellation in the present task.

An important distinction in the inhibitory control literature is between reactive inhibition, the rapid cancellation of an already-initiated response, as measured by the stop-signal reaction time in the stop-signal paradigm, and proactive inhibition. the anticipatory, context-driven suppression of a planned response before it is fully initiated [24]. These two modes of inhibitory control are thought to recruit dissociable neural circuits, Since participants in the present study were informed of the timing of the No-Go cue and could therefore predict its potential arrival, the current protocol likely engaged circuits involved in proactive rather than reactive inhibitory control. This is consistent with the bilateral pre-No-Go beta suppression observed here, which may reflect motor preparation rather than a reactive inhibitory response, and with the absence of a rapid, stimulus-locked beta increase of the kind typically associated with reactive stopping. Proactive inhibition has been specifically associated with effector-specific cortical beta modulation in prior work. The present findings suggest that the effector-specific beta rebound observed here reflects a proactive, targeted suppression of the specific motor representation being cancelled, rather than a global reactive inhibitory broadcast.

An important observation in the present data is the relatively late emergence of the beta rebound, which peaked in the 500–1000 ms window following the No-Go cue at both the cortical and peripheral levels. This timing is later than the rapid beta increases typically reported in reactive stop-signal studies, where beta activity often begins to increase within 200–400 ms of the stop signal [25], and is more consistent with the timescale of the post-movement beta rebound (PMBR), characterised as an increase in beta power that follows voluntary movement termination and is thought to reflect a resetting of the motor system [2]. Under this alternative framework, the bilateral beta suppression observed in the pre-stimulus and early post-No-Go windows may reflect the motor planning process itself, that is, the preparation and partial initiation of the planned wrist movement, rather than a purely inhibitory signal. The subsequent beta rebound would then represent not an inhibition-specific synchronisation, but rather the classic post-planning resetting of the motor system following a period of increased motor cortical activity, analogous to the PMBR observed after completed movements [2], [3]. This interpretation is supported by the bilateral nature of the suppression, which is condition-independent and therefore unlikely to reflect effector-specific inhibitory processing, and by the temporal profile of the rebound, which is more consistent with a PMBR than with a rapid reactive stop signal. Regardless of whether the observed modulation is best characterised as an inhibition-specific beta rebound or a PMBR, the finding that it is effector-specific and dissociable from force output raises the question of what functional role this peripheral beta synchronisation serves.

Inspection of the force time courses revealed a clear small numerical but not statistically significant reduction in wrist force in the period immediately following No-Go cue onset, which could in principle represent a partial, aborted movement. Post-movement beta rebounds are well documented following even brief periods of motor cortical activity [2], [3], and it is therefore conceivable that this transient force perturbation, rather than the inhibitory process per se, drove the peripheral beta rebound observed in the 0.5–1 s window. However, several features of the data argue against this interpretation. For instance, the force perturbation following cue onset was temporally invariant across conditions: the force profile did not differ significantly between NoGo-Left and NoGo-Right at any time window, indicating that any transient force reduction was equivalent regardless of which movement was being cancelled. If this perturbation were responsible for the peripheral beta rebound, one would expect the rebound to be similarly non-lateralised, yet the beta rebound was strongly and selectively lateralised to the channel ipsilateral to the cancelled movement. The lateralised beta rebound therefore cannot be accounted for by a non-specific force perturbation, and is more likely attributed to the effector-specific inhibitory process itself. Nonetheless, future studies employing kinematic recordings alongside EMG would be well placed to quantify the magnitude of any aborted movement initiation more precisely, and to determine whether the amplitude of the peripheral beta rebound scales with the degree of partial movement suppression.

The observation that beta power modulation is detectable not only in sensorimotor cortex but also in wrist EMG raises a fundamental question about the functional significance of peripheral beta synchronisation. Beta oscillations in EMG are thought to reflect the coherent discharge of motor units driven by oscillatory corticospinal input [26], and corticomuscular coherence in the beta band has been extensively documented during sustained isometric contractions [13]. However, the functional consequences of this beta projection remain debated. A candidate function was recently proposed by Borzelli et al. [27], who demonstrated that peripheral beta oscillations may be associated with increased muscle co-contraction and joint stiffness. In their study, peripheral beta activity is implicated in the activation of antagonist muscle pairs, increasing joint impedance without necessarily changing joint torque or displacement. In the context of movement cancellation, this framework offers a compelling account of the present findings. The ipsilateral beta rebound in wrist EMG may reflect a mechanism by which the motor system increases joint stiffness at the to-be-cancelled wrist, thereby physically resisting the execution of the planned movement through co-contraction rather than through a purely neural suppression of motor drive. Under this hypothesis, peripheral beta synchronisation serves as a final effector-level mechanism of movement suppression, complementing the cortical inhibitory signal by increasing the mechanical impedance of the joint.

Several experimental approaches could be used to test this co-contraction hypothesis directly. First, simultaneous recording of agonist and antagonist EMG pairs at the wrist (e.g. flexor carpi radialis and extensor carpi radialis) during the cancellation task would allow direct quantification of co-contraction indices and their relationship to the beta rebound. If the hypothesis is correct, the ipsilateral beta rebound should be accompanied by a simultaneous increase in antagonist co-activation, with the co-contraction index correlating with beta power on a trial-by-trial basis. Second, perturbation-based measures of joint stiffness applied during the cancellation period would provide a test of whether the beta rebound is associated with increased mechanical impedance at the cancelled wrist. Third, transcranial magnetic stimulation could be used to probe corticospinal excitability during the beta rebound period, allowing dissociation of the cortical and peripheral contributions to the observed synchronisation.

A notable feature of the present results is the relative magnitude of beta modulation at the cortical and peripheral levels. Although both the EEG and EMG signals exhibited significant post-No-Go beta modulation, the effect sizes associated with the ipsilateral EMG rebound (*d* = +0.73–+1.34) were consistently larger than those observed at the cortical level (*d* = +0.42–+0.83), and the lateralisation of the rebound was clearly stronger in the peripheral signal (Fig.5). The poorer spatial specificity of the cortical signal is likely attributable, at least in part, to the limitations of scalp EEG. In the context of the present paradigm, volume conduction may have caused activity from the contralateral sensorimotor generator to contaminate the ipsilateral cluster, attenuating the lateralisation. It is plausible that more sophisticated pre-processing methods would have resolved the two conditions more cleanly. However, these observations raise the possibility that peripheral EMG recordings may, in some respects, offer a more targeted window into effector-specific sensorimotor dynamics than scalp EEG. Since each wrist EMG channel records activity from a spatially discrete muscle group with a well-defined corticospinal projection, it is selective for the motor representation of that specific effector. The present results suggest that, for paradigms involving selective suppression of a specific effector, peripheral beta acticity may provide a more sensitive index of effector-level inhibitory dynamics than EEG.

## Methods

### Participants

Fifteen able-bodied participants (25*±*2.5 years old) volunteered in this study. All participants were righthanded, possessed normal or corrected-to-normal vision, and had no history of neurological disorders. This study was approved by the Imperial College Research Ethics Committees (ICREC reference 18IC4685). Prior to the experiment, participants received an explanation of the experiment procedure and provided signed informed consent.

### Experimental protocol

Participants were seated opposite a computer screen with both forearms secured in a custom-built force platform, the dorsal surface of each hand in contact with a dedicated force sensor (Fig. 6a). Two horizontally scrolling force traces were displayed in real time, with vertical displacement proportional to wrist extensor force: red for the left wrist and blue for the right (Fig. 6b). Surface EMG from the left and right wrist extensors and continuous EEG were recorded concurrently throughout the experiment.

**Figure 6.**
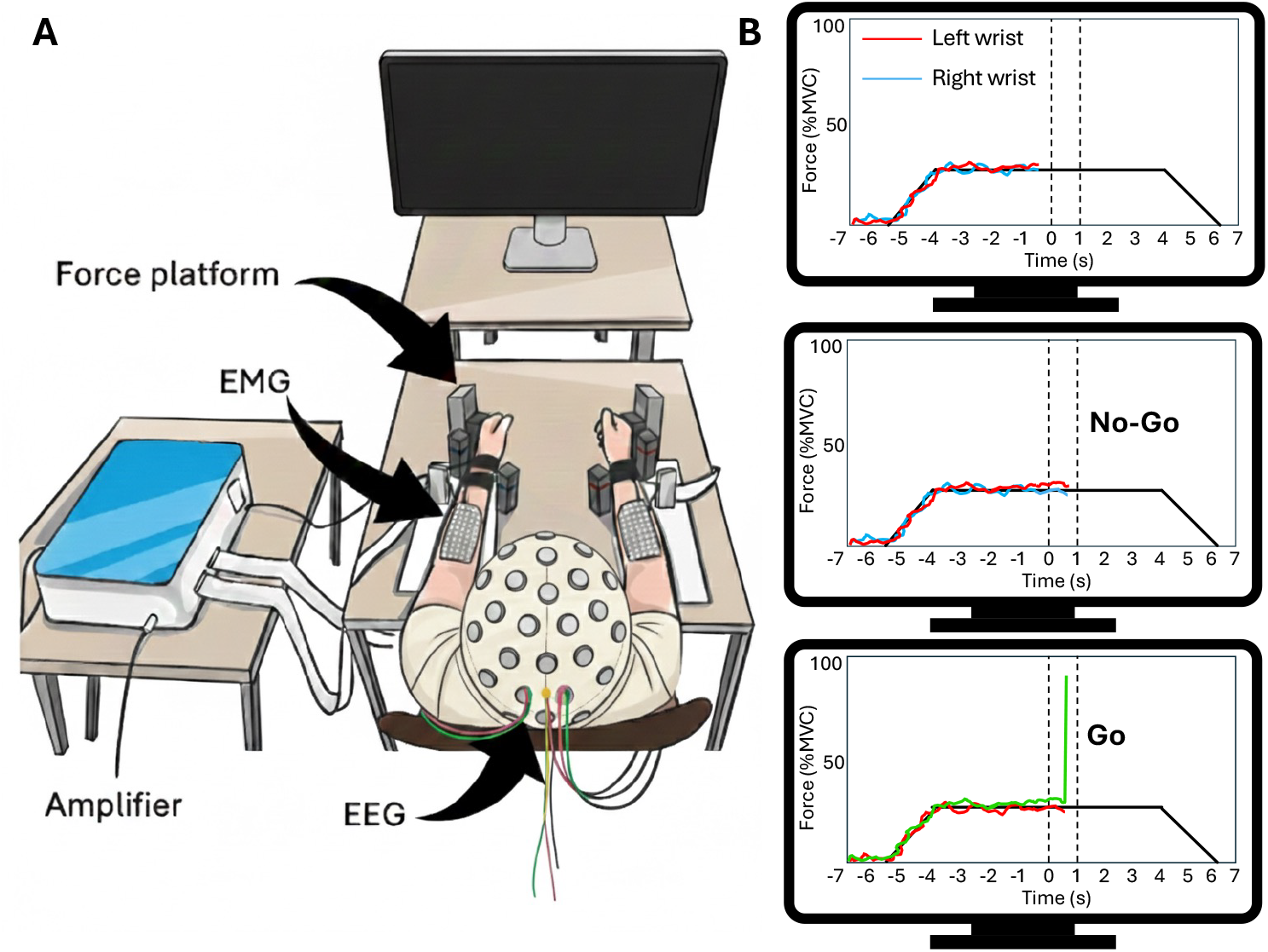
Experimental Setup: **A** Participants viewed Go/No-Go cues and real-time bilateral wrist extensor force while EEG and EMG signals were recorded.**B** Task display showing left and right force traces; middle: No-Go trial, lower: right Go trial.

#### Maximum voluntary contraction

Prior to the main task, each participant’s maximum voluntary contraction (MVC) was determined. Participants performed three 5 s maximal bimanual wrist extensions against the force sensors, with verbal encouragement provided. The peak force recorded across the three attempts was used to scale the force display for the remainder of the session.

#### Task structure

Each trial followed a fixed structure: 1 s rest, a 1 s linear ramp to the target force level (10 % MVC), a 10 s steady-state plateau, a 1 s linear ramp-down, and 1 s rest (14 s total). During the plateau phase, participants maintained bilateral isometric wrist extensor force at 10 % MVC, guided by the real-time force display. Two vertical markers on the display indicated the timing of the Go/No-Go cue, presented at 7 s into the plateau, and, on Go trials, the target response window.

On Go trials (50 % of trials), one force trace turned green at the cue, instructing the participant to perform a rapid, unilateral ballistic wrist extension exceeding 60 % MVC with the pre-specified hand. The responding hand was held constant within each run and alternated across runs. On No-Go trials (50 % of trials), no visual cue was presented and participants maintained steady bilateral force, withholding any movement.

#### Experimental design

The experiment comprised of 10 runs of 20 trials each (10 Go, 10 No-Go per run), yielding 100 trials per condition. The active hand alternated across runs, resulting in five runs per hand. Rest periods of 1–2 min were provided between runs to minimise fatigue. Prior to data collection, all participants completed one practice run to ensure task comprehension.

### Data acquisition

#### EEG acquisition and pre-processing

Cortical activity was recorded from 64 active EEG electrodes (actiCHamp, Brain Products GmbH, Gilching, Germany) at 250 Hz using the international 10-10 system [28]. AFz served as ground and the reference was the average of both earlobes. EEG was high-pass filtered at 0.1 Hz, and a 50 Hz notch filter was applied. Electrode impedance was maintained below 10 kΩ using conductive gel and light abrasion.

EEG preprocessing was performed using MNE-Python [29]. Continuous signals were band-pass filtered (1–100 Hz, 3rd order Butterworth), before being epoched relative to cue onset. Artefact rejection was performed in two stages. Epochs were first visually inspected, and channels exhibiting sustained noise or signal dropout were removed and reconstructed via spherical spline interpolation. Independent Component Analysis (ICA; *n* = 15 components) was then applied to the epoched data, and components attributable to ocular artefacts were removed, as identified by an experienced researcher. A common average reference was applied following artefact rejection.

#### EMG acquisition and pre-processing

Surface EMG signals were recorded from the left and right extensor carpi radialis muscles using highdensity EMG electrode arrays (GR04MM1305, OT Bioelettronica, Turin, Italy), each comprising of 64 monopolar channels arranged in a 13-row*×*5-column configuration, with an inter-electrode distance of 4 mm. Signals were amplified and digitised using a Quattrocento bioelectric signal amplifier (OT Bioelettronica, Turin, Italy) with an internal band-pass filter of 10–500 Hz and sampled at 2,048 Hz. Each electrode grid was referenced to a ground electrode placed around the ipsilateral wrist.

For analysis, a bipolar derivation was computed as the difference between the column-wise means of two spatially separated electrode rows: 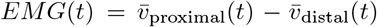 where 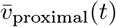 and 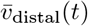 denote the mean monopolar potentials across the proximal and distal reference rows, respectively. This spatial subtraction acts as a high-pass filter along the proximal–distal axis of the muscle, attenuating spatially uniform interference whilst preserving the underlying neuromuscular signal. Bipolar EMG signals were high-pass filtered at 10 Hz and transformed using the absolute value of the Hilbert transform, providing a time-resolved estimate of motor unit activity [30]. EEG and EMG signals were synchronised via a common digital trigger.

#### Force acquisition

Force was recorded using a custom-built force platform equipped with analogue load sensors (FC2231-0000-0100-L; Mouser Electronics, Buckinghamshire, UK) connected to the auxiliary channels of the Quattrocento amplifier (See Fig. 6a). Signals from the left and right wrist extensors were low-pass filtered at 10 Hz (fourth-order, zero-phase) and normalised to each participant’s MVC. Force was used to screen trials for compliance with the No-Go task. Trials were excluded if, within a *±*3 s window centred on the No-Go cue, mean force fell outside the target range (9–11 % MVC) or force variability exceeded 0.5 % MVC (standard deviation).

### Time-frequency analysis

Trials were included in the analysis only if all three data streams met concurrent quality criteria: EEG epochs were free of artefacts as determined by amplitude-based rejection (peak-to-peak threshold *<*70 *µ*V across all EEG channels), EMG peak-to-peak amplitude was below 2 *m*V, and force output complied with the criteria described above. Trials failing any single criterion were excluded in their entirety, ensuring that EEG, EMG and force data were temporally matched across all retained observations.

Cortical and peripheral broadband power (5–50 Hz) was extracted from each trial using Morlet wavelet convolution [31], implemented using the `compute_tfr` function in `MNE-Python` (v1.7) with frequency-adaptive cycle counts (*n*_cycles_ = *f/*2, where *f* is the centre frequency). Trial-level power estimates were normalised using the following *z*-score procedure [32]. First, each trial was standardised relative to its own mean and standard deviation across the full trial, removing trial-level differences in absolute power. Second, the resulting values were re-referenced to a common baseline window (−2 to −1 s pre-NoGo cue), computed by pooling all trials within each subject and normalising by the mean and standard deviation of the baseline distribution. This procedure yields *z*-scored power values that are interpretable as deviations from the pre-NoGo cue baseline in units of standard deviation, while controlling for between-trial variability in signal amplitude [32].

To consider the lateralisation of beta power at the cortical level, two sensorimotor electrode clusters were derived: a left hemisphere cluster comprising of electrodes F3, FC5, FC3, FC1, C5, C3, C1, CP5, CP3, CP1, P1 and P3, and a right hemisphere cluster comprising of F4, FC6, FC4, FC2, C6, C4, C2, CP6, CP4, CP2, P2 and P4. These clusters were defined based on established sensorimotor topographies [33] relating to movement cancellation and to motor activities concerning the wrists and were held constant across all subjects. For each trial, cluster-averaged beta power was computed by averaging the *z*-scored time–frequency representation across electrodes, frequencies (15–35 Hz) and timepoints within each of three pre-specified time windows: a pre-cancellation window (−0.5–0 s), an early cancellation window (0–0.5 s) and a late cancellation window (0.5–1 s). Peripheral beta power was extracted similarly from single-channel EMG recordings at the left and right wrist extensors, applying identical normalisation and time–frequency procedures.

### Statistics

All statistical analyses were performed in Python 3.13 using the `statsmodels` (v0.14) [34], `scipy` (v1.13 [35]), and `pandas` (v2.2) [36] libraries. The significance threshold was set at *α* = 0.05 throughout.

#### Linear mixed model

The primary statistical analysis used a linear mixed model (LMM) [37] fitted by restricted maximum likelihood (REML) [38] using the `MixedLM` implementation in `statsmodels`. The outcome variable was trial-level *z*-scored beta power. Channel (EEG analysis: left sensorimotor cluster, right sensorimotor cluster; EMG analysis: left wrist, right wrist), Condition (NoGo-Left, NoGo-Right) and Time Window (−0.5–0 s, 0–0.5 s, 0.5–1 s) were entered as within-subject fixed effects with a full factorial interaction structure. Subject was modelled as a grouping factor. The random effects structure was determined empirically by comparing nested models using likelihood ratio tests (LRT) on maximum-likelihood (ML) estimates. A model with random intercepts and random slopes for Condition per subject was compared against a random-intercept-only model. The random slope structure was retained if the LRT was significant at *α* = 0.05, indicating meaningful between-subject variability in condition-dependent lateralisation. The inclusion of trial count as an additional fixed-effects covariate was evaluated by a further LRT against the intercept-only model. This term was omitted if it did not significantly improve fit. The selected random effects structure was then refit using REML for final parameter estimation. Fixed effects were evaluated using Type III Wald *χ*^2^ tests. Effect sizes for omnibus fixed effects are reported as partial eta-squared, approximated from the Wald statistic as 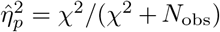, where *N*_obs_ is the total number of trial-level observations. As this approximation is computed relative to the full trial-level sample size, it systematically underestimates effect magnitude at the subject level. Cohen’s *d* from post-hoc comparisons is therefore the primary effect size metric reported.

#### Post-hoc comparisons

Two families of post-hoc comparisons were conducted on subject-level means, computed by averaging trial-level values within each subject, channel, condition and time window cell. The first family comprised pairwise condition contrasts (No-Go-Left vs. No-Go-Right) within each hemisphere and time window combination (*n* = 6 comparisons), tested using paired-samples *t*-tests and corrected using the Holm procedure [39]. The second family comprised one-sample *t*-tests of each channel *×* condition *×* window cell against zero (*n* = 12 comparisons), testing whether beta power was significantly modulated from the baseline. This tests were corrected using the Benjamini–Hochberg false discovery rate (FDR) procedure [40], which is more appropriate than family-wise error rate control for a set of correlated within-subject comparisons. Cohen’s *d* is reported for all post-hoc tests, computed as the mean divided by the standard deviation for one-sample tests, and as the mean difference divided by the standard deviation of the difference scores for paired tests.

#### Cluster-based permutation

To visualise periods of significant beta-band power modulation relative to baseline, we applied a non-parametric cluster-based permutation test [41] using MNE-Python [29]. At each time point, a one-sample *t*-statistic was computed across subjects testing against a null hypothesis of no modulation. Contiguous time points exceeding a cluster-forming threshold of *t*^∗^ = *t*_0.975,*ν*_, where *ν* = *n* − 1, were grouped into candidate clusters, with cluster-level statistics defined as the sum of constituent *t*-values. Significance was assessed by comparing observed cluster statistics against a null distribution of maximum cluster statistics derived from *B* = 5,000 sign-flipping permutations. Clusters were considered statistically significant at *p <* 0.05.

## Acknowledgments

This study was supported by UK Research and Innovation (UKRI) under the UK Government’s Horizon Europe funding scheme (Grant number: 10052152; HybridNeuro).

## Notes

### Competing Interest Statement

The authors have declared no competing interest.

